# Genetic inactivation of the translin/trax microRNA-degrading enzyme phenocopies the robust adiposity induced by *Translin* (*Tsn*) deletion

**DOI:** 10.1101/2020.04.25.060665

**Authors:** Xiuping Fu, Aparna P. Shah, Zhi Li, Mengni Li, Kellie L. Tamashiro, Jay M. Baraban

## Abstract

**Objective:** Deletion of *Translin* (*Tsn*) from mice induces an unusual metabolic profile characterized by robust adiposity, normal body weight and glucose tolerance. Translin (TN) protein and its partner, trax (TX), form the TN/TX microRNA-degrading enzyme. Since the microRNA system plays a prominent role in regulating metabolism, we reasoned that the metabolic profile displayed by *Tsn* KO mice might reflect dysregulation of microRNA signaling.

**Methods:** To test this hypothesis, we inserted a mutation, E126A, in *Tsnax*, the gene encoding TX, that abolishes the microRNA-degrading enzymatic activity of the TN/TX complex. In addition, to help define the cell types that drive the adiposity phenotype, we have also generated mice with floxed alleles of Tsn or Tsnax.

**Results:** Introduction of the E126A mutation in *Tsnax* does not impair expression of TN or TX proteins or their co-precipitation. Furthermore, these mice display selective increases in microRNAs that match those induced by *Tsn* deletion, confirming that this mutation in *Tsnax* inactivates the microRNA-degrading activity of the TN/TX complex. Mice homozygous for the *Tsnax* (E126A) mutation display a metabolic profile that closely mimics that of *Tsn* KO mice.

Selective deletion of *Tsn* or *Tsnax* from either adipocytes or hepatocytes, two candidate cell types, does not phenocopy the elevated adiposity displayed by mice with constitutive *Tsn* deletion or the *Tsnax*(E126A) mutation. Furthermore, global, conditional deletion of *Tsn* in adulthood does not elicit increased adiposity.

**Conclusion:** Taken together, these findings indicate that inactivation of the TN/TX microRNA-degrading enzyme during development is necessary to drive the robust adiposity displayed by *Tsn* KO mice.

## 1. Introduction

Adipose tissue plays a central role in orchestrating metabolic homeostasis and the pathophysiology of metabolic disorders. Therefore, it is imperative to decipher the molecular mechanisms that control its size and health [1–3]. Analysis of mice carrying genetic mutations that elicit abnormalities in adipose tissue has yielded important insights into these processes [4]. Accordingly, we have in recent studies characterized the metabolic profile of *Tsn* KO mice that display robust adiposity in the context of normal body weight and found that these mice retain normal glucose tolerance [5]. Those findings have heightened interest in identifying the molecular mechanism linking Tsn deletion to adiposity.

Initial characterization of translin (TN) protein revealed that it shares homology and forms a complex with TN-associated protein X, or trax (TX) [6, 7]. Furthermore, deletion of Tsn in mouse, Drosophila or yeast leads to loss of TX protein, suggesting that the stability of TX is dependent on its physical interaction with TN [8]. A major breakthrough in understanding the function of the TN/TX complex emerged from Drosophila studies that demonstrated that it possesses RNase activity and mediates processing of microRNAs [9]. Subsequent studies in mouse revealed that this complex acts as a microRNA-degrading enzyme which targets a small subpopulation of microRNAs *in vivo* [10, 11]. For example, examination of the impact of *Tsn* deletion on microRNA profiles in cerebellum, hippocampus and aorta have identified small, partially overlapping cohorts of microRNAs that are elevated in each of these tissues[10, 12, 13].

Recent studies have strongly implicated the microRNA system in regulating adipose tissue size and function [14–16]. For example, conditional deletion of Dicer from adipocytes inhibits lipogenesis in white adipocytes and produces severe depletion of white adipose tissue [17, 18]. In previous studies, we have shown that the TN/TX complex can oppose the action of Dicer by degrading pre-microRNAs, thereby preventing their processing into mature microRNAs by Dicer [10]. Thus, these findings suggested that the adiposity displayed by *Tsn* KO mice could be attributed to increased microRNA signaling due to loss of the TN/TX microRNA-degrading enzyme. To test this hypothesis directly, we have taken advantage of recent studies which have demonstrated that a point mutation in *Tsnax*, E126A, abolishes the RNase activity of this enzyme [9, 10]. Accordingly, we generated mice with this point mutation in *Tsnax* and investigated if this point mutation is sufficient to phenocopy the adiposity and metabolic profile displayed by *Tsn* KO mice.

## 2. Materials and Methods

### 2.1. Mice

All experimental procedures were performed in accordance with the NIH’s Guide for the Care and Use of Laboratory Animals and approved by the Johns Hopkins Animal Care and Use Committee. A colony of *Tsn* KO mice was established at Johns Hopkins University from the line generated in Dr. Kasai’s laboratory [19] and provided by the JCRB Laboratory Animal Resource Bank of the National Institute of Biomedical Innovation (*Tsn* KO: Nbio055). These mice had been backcrossed to C57BL/6 mice for over ten generations. Mice were housed in ventilated racks, on a 14-h/10-h light/dark cycle and with standard chow (2018SX Teklad Global, Frederick, MD; unless stated otherwise) and free access to tap water.

### 2.2. Generation of mice with the E126A point mutation in Tsnax

Mice containing the E126A mutation in the Trax coding region were generated on a C57 background based on a one-step procedure for generation of mutations in mice using CRISPR/Cas9 technology [20]. Briefly, we used the MIT guide RNA design tool to select a sgRNA (5’-TTTAGGACTGCAGGAATACG-3’). A single-stranded donor oligo containing the mutant nucleotide sequence was synthesized with ∼30 bp homology arms flanking the predicted DSB (double-strand break) site.

Pronuclear injection of one-cell C57BL/6J embryos (Jackson Laboratories) was performed by the JHU Transgenic Core using standard microinjection techniques [21] with a mix of Cas9 protein (30ng/ul, PNABio), tracrRNA (0.6µM, IDT), crRNA (0.6µM, IDT) and ssDNA oligo (12ng/ul, IDT) diluted in RNAse free injection buffer (10 mM Tris-HCl, pH 7.4, 0.25 mM EDTA). Injected embryos were then transferred into the oviducts of pseudopregnant ICR females (Envigo). Three weeks later, 31 pups were born and genotyped by using the following primers: TraxE126A-F1: 5-AACACAGTCTGCATGGCATC-3, TraxE126A-R1: 5’-CCACCAATCATTATGCTGCTG-3’ to generate PCR products, which were then incubated with the StuI restriction enzyme. As the mutant sequence introduces a StuI restriction site, cleavage by StuI indicates successful insertion of the mutant sequence. Using this assay, we identified 5 mice containing a mutant allele. This inference was confirmed by sequencing of the PCR products. One of these mice was used to establish the colony. In addition, we amplified and sequenced the top 5 off-target sites as predicted by the MIT guide RNA design tool. No off-target genomic alterations were found.

### 2.3. Generation of mice with floxed alleles of Tsn or Tsnax

Mice with a floxed allele of *Tsn* were also generated on a C57 background by using the easi-CRISPR protocol [22]. We designed one sgRNA (5’ –TTATCCGTCCTATTGCTAGA-3’) targeting *Tsn* intron 1 and one sgRNA (5’-ATAGGGGTTTGGTCATTTTG-3’) targeting *Tsn* intron 2. A long single-stranded donor oligo was synthesized that spanned exon 1 and also contained two loxP sites as well as 66 bp homology arms that match segments flanking the predicted DSB sites. One-cell C57BL/6J embryos were pronuclear injected and transferred to the oviducts of pseudopregnant ICR females as described above. Seven pups were born and genotyped by PCR using the following primers: TSN-F1: 5’-TGACCTCGAACTCGAACCTGT-3’, LoxP-R: 5’-CGTATAATGTATGCTATACGAAG-3’. One of these mice contained the correct insertion of loxP sites flanking exon 1 of *Tsn* as confirmed by sequencing. This animal was used to establish the colony. We amplified and sequenced the top 10 off-target sites based on MIT guide RNA design tool. No off-target genomic alterations were detected.

Mice with a floxed allele of *Tsnax* were also generated on a C57 background by CRISPR/Cas9 technology. We designed one sgRNA (5’-TGTGCTAGCGCGGCATCGCA-3’) targeting *Tsnax* intron 1, one sgRNA (5’-TGCGGTGGCTTAGCGAGTAA-3’) targeting *Tsnax* intron 3, along with two single-stranded donor oligos that contained a single loxP site and different flanking homology arms for each of the two DSB sites. One-cell C57BL/6J embryos were pronuclear injected and transferred to the oviducts of pseudopregnant ICR females as described above. Twenty-four pups were born and genotyped by PCR using the following primers: TX F1: 5’-ACCTGTGTGTGGCTGGAGA-3’, TX F2: 5’-ATGTGTTCTTCCTGTCG-3’, LoxP-R: 5’-CGTATAATGTATGCTATACGAAG3’. Only one mouse had the correct insertion of one of the distal loxP site. This animal was used to generate a colony of mice that were homozygous or heterozygous for insertion of the distal loxP site. Oocytes harvested from these mice were used for a microinjection of the sgRNA and donor oligo targeting intron1. Thirteen pups were born and genotyped. Only one mouse was found to have an allele that contained both loxP sites as determined by sequencing. This animal was used to establish the colony of *Tsnax* conditional KO mice. The top 10 off-target sites predicted by the MIT guide RNA design tool were screened and no evidence of off-target genomic alterations was found. All sgRNAs and single-stranded oligos were purchased from IDT (Coralville, IA).

To generate mice with conditional deletion of either *Tsn* or *Tsnax*, we bred mice with one of these floxed alleles with mice from one of the following Cre lines obtained from Jackson Labs: Adipoq-Cre (#028020), Alb-Cre (#003574), or Ubiquitin C-CreERT2 (# 007001). All conditional KO mice used for this study were hemizygous for the Cre allele and homozygous for the floxed allele. Mouse genotyping was routinely performed on tail snips by Transnetyx, Inc. (Cordova, TN).

### 2.4. Tamoxifen Treatment

Tamoxifen (MilliporeSigma, Burlington, MA) solution (10mg/ml in corn oil) was administered to mice with the ubiquitin C-CreERT2 allele by intraperitoneal (i.p.) injection at a dose of 100 mg/kg once daily for 6 consecutive days.

### 2.5. Body composition, length and muscle mass

Body composition was determined by using a nuclear magnetic resonance scanner (EchoMRI-100, Houston, TX). Body length was measured as the distance from the tip of the snout to the base of the tail. The weight of the quadriceps femoris muscle was used as a gauge of muscle mass. Determination of subcutaneous and visceral fat depots was performed as described in Shah et al [5].

### 2.6. Glucose Tolerance Test

To perform the glucose tolerance test (GTT), mice were fasted overnight (∼16 h) and then habituated to the testing room for 1 h. Baseline glucose levels were measured in blood obtained by tail nick using handheld glucometers (Freestyle; Abbott, Alameda, CA, USA). In addition, blood was collected in heparin-coated capillary tubes for analysis of baseline plasma insulin levels. Mice were then injected intraperitoneally with a 20% glucose solution at a dose of 1.5 mg/g body weight. Blood glucose level measurements and blood collection for insulin levels occurred at 15, 30, 45, 60 and 120 min after glucose injection. Blood was centrifuged at 4°C at 3000 RPM for 20 min to obtain plasma for assessing insulin levels by ELISA (described below).

### 2.7. Tissue collection

Mice were fasted for about 6 h at the beginning of their light cycle and killed by rapid decapitation. Blood was collected into EDTA coated collection tubes. Blood was centrifuged at 4°C at 3000 x g for 20 min and plasma was collected and stored at -80°C until analysis by ELISA. Brain samples were collected and frozen immediately on powdered dry ice and stored at -80°C until analysis by RT-PCR or immunoblot. Liver and epididymal white adipose tissue (eWAT) were dissected, flash frozen in liquid nitrogen, and stored at -80°C until analysis by ELISA, RT-PCR, or immunoblot.

### 2.8. Plasma analysis

Plasma adiponectin and leptin were measured by ELISA (#EZMADP-60K and EZML-82K, respectively, MilliporeSigma, Billerica, MA). Plasma FFAs were measured using fluorometry (#700310, Cayman Chemicals, Ann Arbor, MI). Plasma and liver triglycerides were measured using colorimetry (#10010303, Cayman Chemicals). For samples collected during the GTT, plasma insulin levels were measured by ELISA (#90080, Crystal Chem, Downers Grove, IL).

### 2.9. Real-Time Quantitative RT-PCR

Total RNA was isolated from cerebellum, eWAT and liver using the miRNeasy kit (Qiagen, Hilden, Germany). For gene expression analysis, cDNA was synthesized from 200 ng of total RNA with the High-Capacity cDNA Synthesis kit (Applied Biosystems, Foster City, CA). qRT-PCR was performed using the 2x Fast SYBR Green Master Mix (Applied Biosystems). MicroRNAs were reverse-transcribed with specific Taqman primers (Applied Biosystems), and subsequently measured by real-time PCR with the Taqman universal PCR master mix and specific Taqman probes (Applied Biosystems). Sequences of primers are shown in Supplementary Table 2.

### 2.10. Co-Immunoprecipitation (Co-IP)

Pierce Classic IP Kit (Thermo Fisher Scientific, Waltham, MA) was used for Co-IP experiments following the manufacturer’s instructions. Briefly, forebrain samples were homogenized with lysis buffer and protein concentrations determined using the Pierce BCA Protein Assay Kit (Thermo Fisher Scientific). Equal amounts of total protein were incubated with TX antibodies (generated in our lab[23]) overnight at 4°C and then loaded on Pierce Protein A/G Agarose resin for another 1h. After washing 3 times with Wash buffer, followed by an additional wash with Conditioning buffer, samples were heated at 100°C in Laemmli buffer (Bio-Rad, Hercules, CA) and processed for immunoblotting.

### 2.11. Immunoblotting

Forebrain and liver tissue samples were homogenized with RIPA buffer (Cell Signaling Technology, Danvers, MA) containing a cocktail of protease and phosphatase inhibitors (MilliporeSigma, Burlington, MA). Total protein was extracted from eWAT using the Minute Total Protein Extraction Kit for Adipose Tissues (Invent Biotechnologies, Plymouth, MN) following the manufacturer’s instructions. Equal amounts of total protein were separated by an SDS-PAGE gel, transferred to a PVDF membrane (Bio-Rad) and immunoblotted with antibodies: anti-HSC70 (MilliporeSigma, Burlington, MA), anti-actin (MilliporeSigma) and anti-tubulin (Cell Signaling Technology), anti-TN and anti-TX (generated in our lab). Blots were developed with the ECL system (Thermo Fisher Scientific). Band intensities were quantified from digital images by densitometry using ImageJ.

### 2.12. Histology

5-µm-thick sections of paraffin-embedded liver samples were stained with haematoxylin and eosin (H&E) and imaged using bright field microscopy (KEYENCE BZ-X700, Itasca, IL).

### 2.13. TN/TX RNase assay

A synthetic miR-409-5p oligo was 5’ end-labelled with γ-P^32^ ATP using T4 polynucleotide kinase (New England Biolabs) and purified using G-50 columns. The labelled oligo was annealed with unlabelled synthetic miR-409-3p by heating the mixture to 65 degrees C and then allowing it to cool slowly to room temperature. Recombinant wild type TN/TX and mutant TN/TX(E126A) complexes were prepared as described in Asada et al.[10] and provided by Z. Paroo. Cleavage assays were conducted in 100mM KCl, 20mM Tris with either 3mM MgCl_2_ or 2.5mM EDTA at 37 degrees C for 15 minutes. Following incubation, the reactions were immediately loaded on to 4.5% native polyacrylamide gels.

### 2.14. Statistical Analysis

Data are presented as mean ± SEM. Statistical significance was evaluated using GraphPad Prism 7 (GraphPad Software, La Jolla, CA). Student’s t-test or one-way ANOVA were used to compare groups in single-variable experiments. Ordinary or repeated measures (RM) two-way ANOVA were used to analyze multiple-variable experiments. Pairwise comparisons were made using Bonferroni’s post-hoc test. Differences were considered significant at P < 0.05.

## 3. Results

### 3.1. Generation and initial characterization of Tsnax(E126A) mice

Prior to introducing the E126A mutation into the mouse *Tsnax* coding region, we first confirmed that this mutation abolishes the activity of the TN/TX microRNA-degrading enzyme. Accordingly, we compared the ability of recombinant TN/TX complexes generated with wild type (WT) TN and either WT or mutant (E126A) TX to cleave miR-409 in vitro. miR-409 was selected as an in vitro substrate because we have previously shown that this microRNA is elevated in cerebellum and hippocampus of TN KO mice [10]. As expected, the WT TN/TX complex produced near complete cleavage of this substrate, while the mutant TN/TX(E126A) complex did not, indicating that it was inactive (Figure 1A).

**Figure 1:**
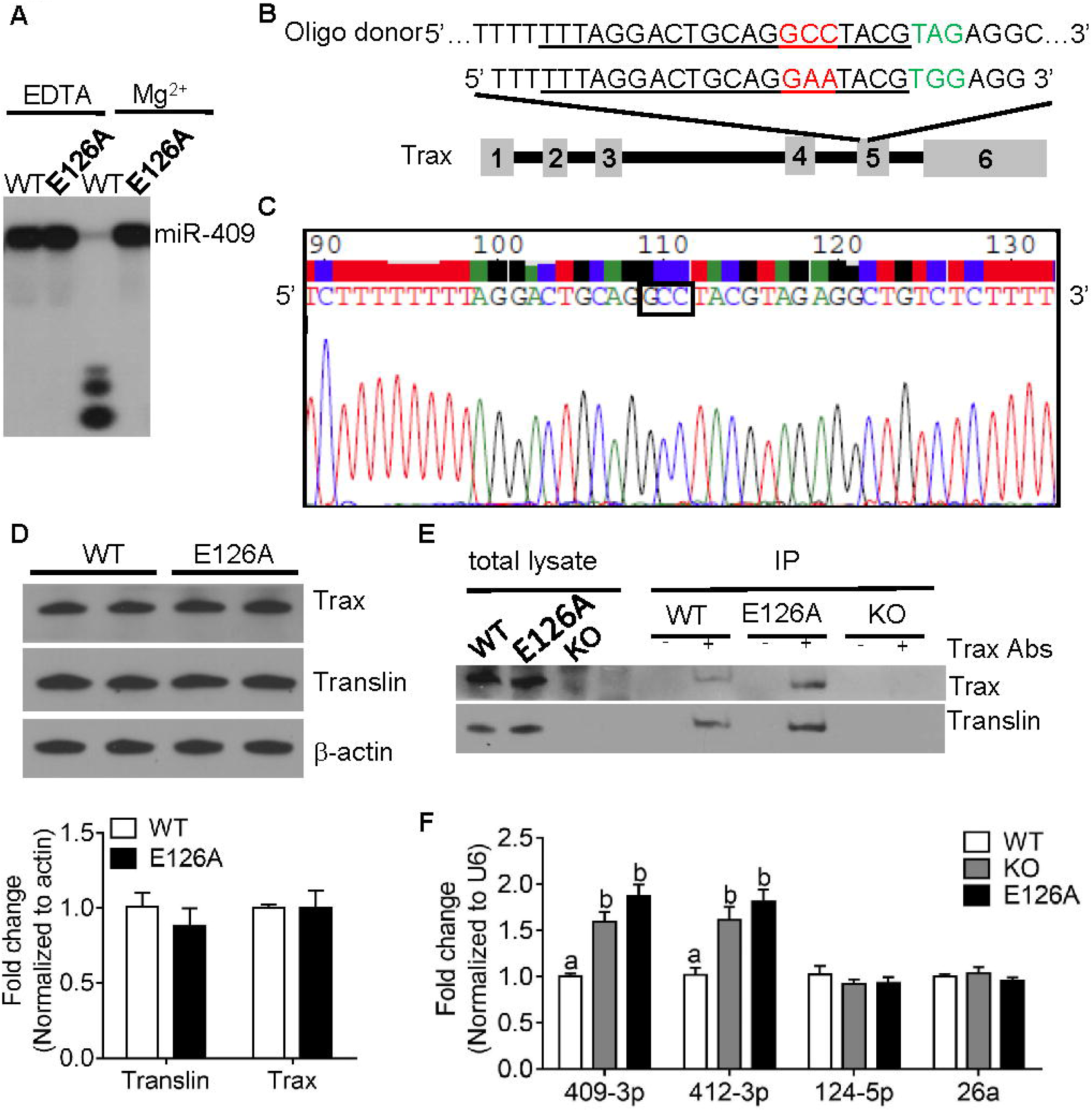
Generation and characterization of TX(E126A) mice. (A) The *Tsnax*(E126A) mutation abolishes the microRNA-degrading activity of the TN/TX complex *in vitro*. Purified recombinant TN/TX complexes were prepared from the native sequence of mouse *Tsn* and either the native mouse *Tsnax* sequence, (WT), or the *Tsnax*(E126A) mutant version. These recombinant complexes were incubated with radiolabeled, double-stranded, mature miR-409. The left two lanes show intact, double-stranded miR-409, as TN/TX RNase activity, which is mg^2+^-dependent [9], is completely blocked by the presence of EDTA. The two right lanes show that the WT TN/TX complex produces near complete cleavage of miR-409 in the presence of mg^2+^. In contrast, the TN/TX complex containing the mutant TX protein is completely inactive in this assay. (B) Schematic representation of the *Tsnax* gene region matching the designed sgRNA and oligo donor for CRISPR. Point mutation is shown in red. PAM site is marked in green. The guide RNA segment is underlined. (C) The nucleotide sequence of the region targeted by the sgRNA confirms successful insertion of the desired mutation. (D) TN and TX protein levels are normal in forebrain samples obtained from *Tsnax*(E126A) mice. n=6/group. Data are expressed as mean ± SEM. Student’s t-test was used for statistical analysis. (E) Co-immunoprecipitation assay shows that the ability of native TN protein to interact with TX is not impaired by insertion of the E126A mutation in TX. (F) MicroRNA profiling in cerebellum. *Tsnax*(E126A) mice display comparable increases in two miRNAs, miR-409-3p and miR-412-3p, that are elevated in *Tsn* KO mice. Two additional miRNAs that are unaffected in *Tsn* KO mice are also unchanged in *Tsnax*(E126A) mice. n=6/group. Data are expressed as mean ± SEM. Different letters indicate statistically significant differences (p < 0.05) according to one-way ANOVA followed by Bonferroni’s post-hoc test.

As these pilot studies indicated that the E126A mutant version of TX abolishes the microRNA-degrading activity of the TN/TX complex, we used the CRISPR/Cas9 approach to introduce this point mutation into the fifth exon of *Tsnax* (Figures 1B and 1C). After generating mice that are homozygous for the mutant allele, we confirmed that the expression levels of both TX and TN proteins are unaffected (Figure 1D). Furthermore, we found that the ability of these proteins to co-precipitate is preserved (Figure 1E).

To compare the effect of introducing this point mutation in *Tsnax* with that of *Tsn* deletion on microRNA levels in cerebellum, we checked the impact of these genetic manipulations on the levels of two microRNAs, miR-409-3p and miR-412-3p, that are elevated in *Tsn* KO mice [10] and of two microRNAs, miR-124-5p and miR-26a that are not. We found that the *Tsnax*(E126A) and *Tsn* KO mice show an identical pattern (Figure 1F) providing compelling evidence that the *Tsnax*(E126A) point mutation inactivates the TN/TX microRNA-degrading enzyme *in vivo*.

### 3.2. Comparison of metabolic profiles of Tsnax(E126A) mice and Tsn KO mice

Following initial characterization of the *Tsnax*(E126A) mice, we proceeded to compare the metabolic profiles of male and female mice that are homozygous for the *Tsnax*(E126A) point mutation or the *Tsn* KO allele. Body composition scans of these mice at 3 months and 6 months of age demonstrated that *Tsnax*(E126A) mice phenocopy the robust adiposity displayed by male and female *Tsn* KO mice, with little effect on body weight (Figures 2A, 2B and Supplementary Figures 1A and 1B). We also found that body length (Figures 2A and 2B) and muscle mass (Supplementary Figure 1C) are decreased to a similar extent in both *Tsn* KO and *Tsnax*(E126A) mice. We previously showed that the elevated adiposity displayed by *Tsn* KO mice reflect increases in both visceral and subcutaneous fat depots, with a small but significant preference for subcutaneous fat[5]. Overall, *Tsn* KO and *Tsnax*(E126A) mice show very similar increases in both visceral and subcutaneous fat depots. However, in calculating the size of these depots as a percentage of total body fat, we noticed a small but significant increase in subcutaneous fat in male, but not in female, *Tsnax*(E126A) mice (Figures 2 C-F).

**Figure 2:**
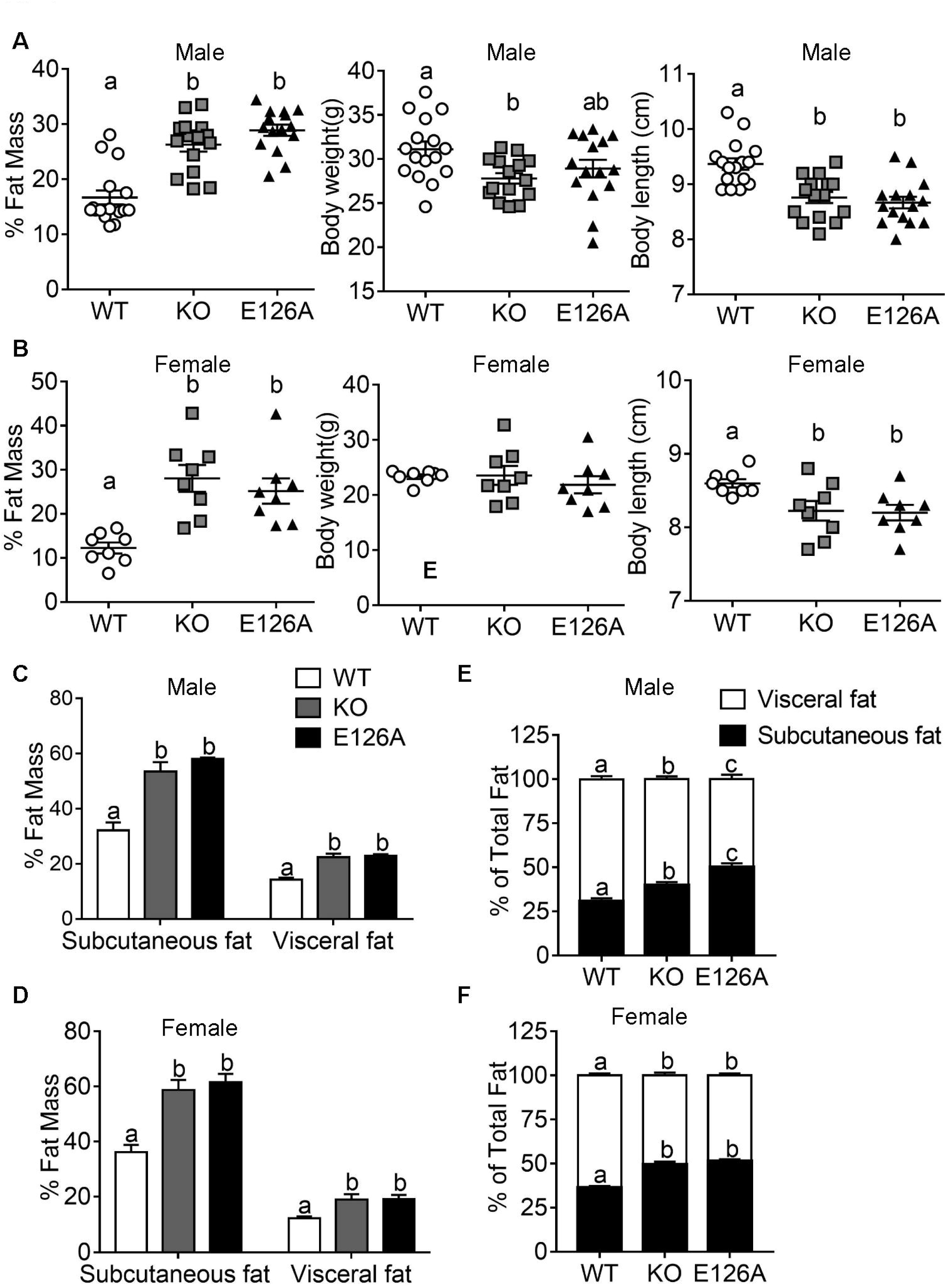
*Tsnax*(E126A) mice phenocopy the increased adiposity displayed by *Ts*n KO mice. Both male (A) and female (B) *Tsnax*(E126A) mice phenocopy the increased adiposity displayed by *Tsn* KO mice at 6 months of age. Male, but not female, *Tsn* KO mice show a significant reduction in body weight that does not reach statistical significance in *Tsnax*(E126A) mice. *Tsn* KO and *Tsnax*(E126A) mice show comparable decreases in body length in both males and females. Both male (C) and female (D) *Tsnax*(E126A) mice show the same pattern of increased subcutaneous and visceral adipose tissue (as percent of carcass weight) displayed by *Tsn* KO mice. (E) Male *Tsnax*(E126A) mice have a greater proportion of subcutaneous than *Tsn* KO mice, while females (F) show the same pattern as *Tsn* KO mice. Males: n=15/group, female: n=8/group. Data are expressed as mean ± SEM. Different letters indicate statistically significant differences (p < 0.05) according to one-way ANOVA followed by Bonferroni’s post-hoc test.

In our previous characterization of the metabolic profile of male *Tsn* KO mice, we unexpectedly found that they retain glucose tolerance despite their robust adiposity[5]. Accordingly, in the present study, we checked if *Tsnax*(E126A) mice, which mimic the adiposity phenotype of *Tsn* KO mice, also retain glucose tolerance. Glucose tolerance testing of male *Tsnax*(E126A) mice demonstrated that their glucose and insulin levels are are not significantly different from those of male *Tsn* KO mice in response to a glucose challenge (Figure 3A and B). In contrast to male *Tsn* KO mice, female *Tsn* KO mice develop significantly higher glucose levels in response to a glucose challenge than WT mice. However, female *Tsn* KO and *Tsnax*(E126A) mice do not differ significantly from each other (Figure 3C and D).

**Figure 3:**
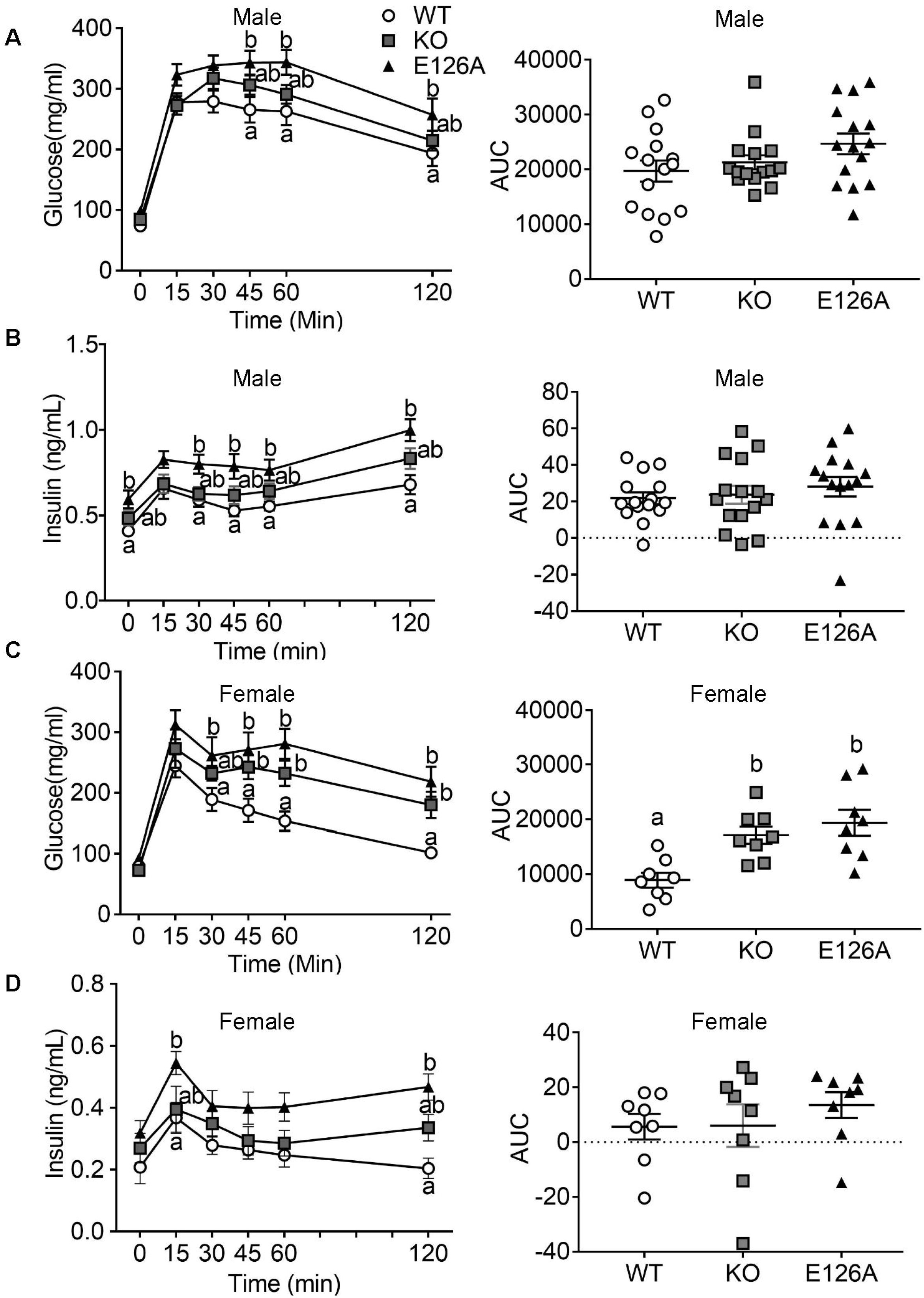
Comparison of Glucose Tolerance Testing in *Tsn* KO and *Tsnax*(E126A) mice. (A) Blood glucose and (B) plasma insulin levels during ip-GTT (1.5 mg/g glucose) of male *Tsn* KO mice are not significantly different from those elicited in male WT mice. Both the glucose and insulin responses to this glucose challenge are significantly higher in *Tsnax*(E126A) male than WT mice, but do not differ from those of *Tsn* KO mice. AUC (area under the curve) of both glucose and insulin are normal in *Tsnax*E126A male mice. n=15/group. (C) Both *Tsn* KO and *Tsnax*(E126A) female mice display elevated blood glucose levels during ip-GTT (1.5 mg/g glucose), compared to WT mice. (D) *Tsnax*(E126A) female mice reach higher plasma insulin levels than WT mice at the 15’ and 120’ time points. n=8/group. Data are expressed as mean ± SEM. Different letters indicate statistically significant differences (p < 0.05) by two-way ANOVA with repeated measures or one-way ANOVA (for AUC) followed by Bonferroni’s post-hoc test.

In our previous study, we identified several candidate protective factors in *Tsn* KO mice that may contribute to their ability to retain glucose tolerance[5]. As described previously and as shown in the current study, these mice have a higher proportion of subcutaneous fat, which is thought to protect against glucose intolerance[24]. Furthermore, *Tsn* KO mice display elevated levels of adiponectin, which correlate with improved glucose tolerance[25, 26]. Consistent with these findings, we found that *Tsn* KO and *Tsnax*(E126A) mice show comparable elevations in plasma adiponectin levels (Figure 4A). Plasma leptin levels are also elevated to a similar extent in these mice (Figure 4B), consistent with their comparable increase in adiposity. Plasma FFAs and TGs are elevated in *Tsn* KO mice compared to WT mice, while *Tsnax*(E126A) mice display values that are not significantly different from either group (Figures 4C and 4D).

**Figure 4:**
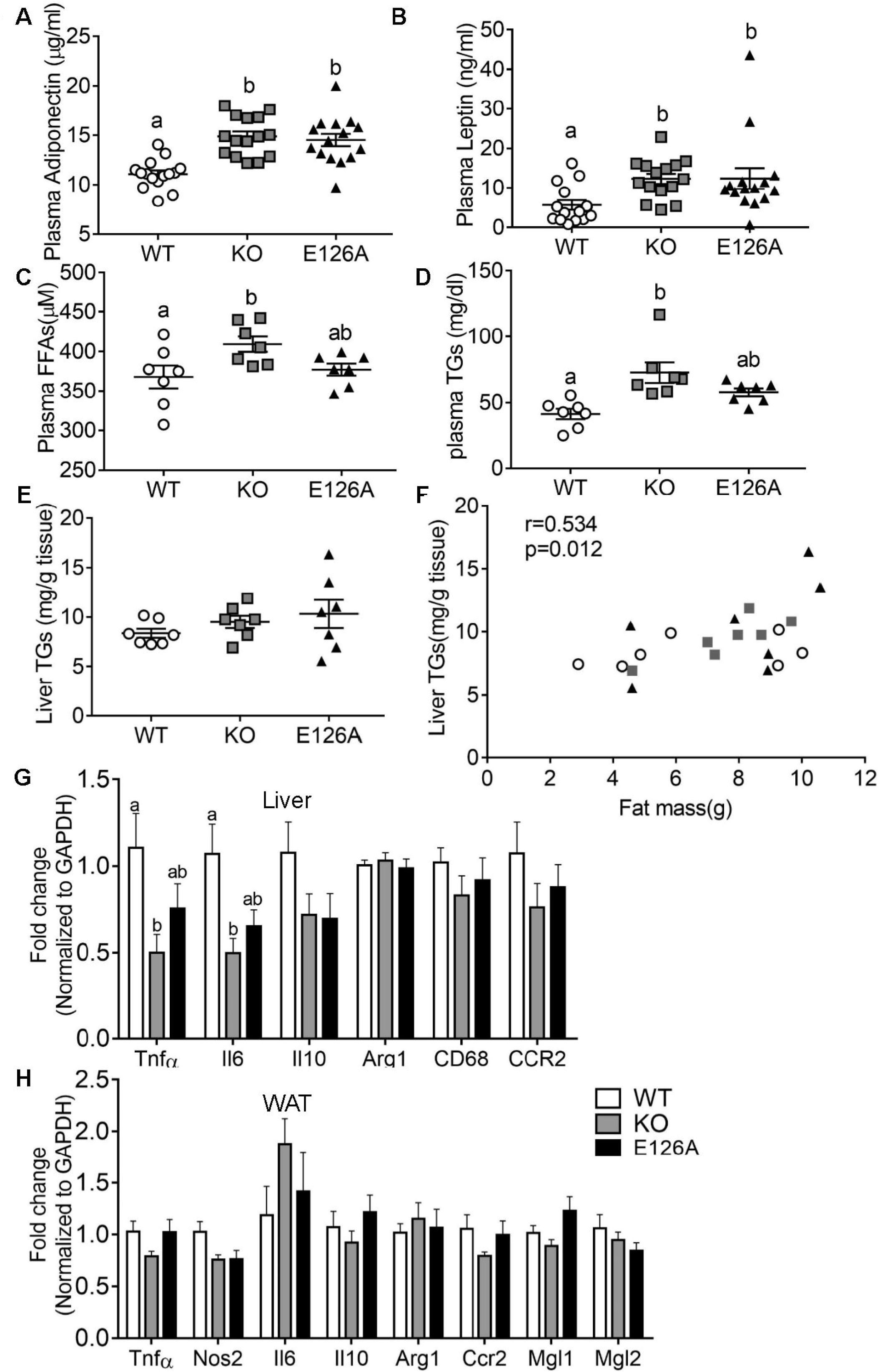
Comparison of metabolic and inflammation markers in *Tsn* KO and *Tsnax*(E126A) mice. (A and B) *Tsnax*(E126A) mice phenocopied the increased plasma levels of adiponectin (A) and leptin (B) found in *Tsn* KO mice. (C and D) TN KO mice have significantly higher plasma free fatty acids (FFAs) and triglycerides (TGs) compared to WT mice, whereas the levels displayed by *Tsnax*(E126A) mice are not significantly different from either of those groups of mice. (E) The average level of liver TGs does not differ across these three groups, however, liver TGs levels correlate with fat mass across these groups (F). (G) Evaluation of the mRNA levels of a panel of inflammation markers in liver shows that expression of TNFα and IL-6 is significantly decreased in *Tsn* KO mice, but not in *Tsnax*(E126A) mice. (H) mRNA levels of inflammation markers in WAT are unaffected in *Tsn* KO and *Tsnax*(E126A) mice compared to WT mice. n=7/group. Data are expressed as mean ± SEM. Different letters indicate statistically significant differences (P < 0.05) by one-way ANOVA followed by Bonferroni’s post-hoc test.

To check if *Tsnax*(E126A) mice also display hepatic steatosis as reported for *Tsn* KO mice[5], we assessed their liver histology. We found that only a subset of both *Tsn* KO mice and *Tsnax*(E126A) mice that had higher levels of adiposity and body weight displayed lipid droplets characteristic of steatosis (Figure S2). Furthermore, we found a correlation between liver TG levels and total fat mass across these groups, even though we did not detect differences between the group means (Figures 4E and 4F). One important variable that might explain the reduced degree of hepatic steatosis observed in Tsn KO mice in this study compared to that seen in our previous report [5] is the use of different diets with distinct composition of carbohydrates and fats. In our previous study, we were comparing the metabolic profile of Tsn KO mice with that of adiposity-matched WT mice. To that end, WT mice were placed on a high-fat diet (D12492, Research Diets Inc.) and Tsn KO mice were fed a matched low-fat diet (D12450J, Research Diets, Inc.). However, in this study, Tsn KO and Tsnax(E126A) mice were fed standard lab chow (2018SX Teklad Global, Envigo). Since the diet used in this study has higher fat content (18%) than that used in the previous study (10%), we cannot attribute the reduced hepatic steatosis observed in this study to reduced fat content of the diet. However, it might be due to other factors, such as differences in the types of fat or the higher levels of carbohydrate content in the previous study.

Another candidate factor that might help *Tsn* KO mice retain glucose tolerance is their reduced level of inflammation in hepatic and adipose tissues [5]. In that study, *Tsn* KO mice show reduced TNFα expression in hepatic tissue compared to WT mice. Comparing TNFα and IL6 mRNA levels in liver samples of *Tsn* KO and *Tsnax*(E126A) mice with those of WT mice confirmed a decrease in these inflammatory markers between WT and TN KO mice. However, the decreases observed in these markers in liver samples from *Tsnax*(E126A) mice were not as large and did not differ significantly from either the WT or *Tsn* KO group (Figure 4G). Consistent with our previous results [5], the expression of inflammatory markers in adipose tissue does not differ among WT, *Tsn* KO and *Tsnax*(E126A) mice (Figure 4H).

### 3.3. Impact of conditional deletion of Tsn or Tsnax from adipocytes on adiposity

While these studies provide compelling evidence that the robust adiposity phenotype displayed by *Tsn* KO mice is due to loss of TN/TX microRNA-degrading activity, we also sought to gain insight into which cell types drive this phenotype. To investigate this question, we generated mice with floxed alleles of *Tsn* or *Tsnax* (Figures S3 and S4) and then examined the effect of selective deletion of these genes in two candidate cell types, adipocytes and hepatocytes.

To assess whether deletion of *Tsn* or *Tsnax* from adipocytes is sufficient to mimic the robust adiposity phenotype, we generated mice that are homozygous for the floxed alleles of TN or TX and hemizygous for the adiponectin-Cre allele, which drives Cre expression selectively in adipocytes[27]. First, we checked that the loxP sites flanking exon 2 of either *Tsn* or *Tsnax* are functional by confirming that TN and/or TX protein expression is reduced selectively in adipose tissue, but not brain, as expected (Figures 5A and 5D). Of note, conditional deletion of *Tsn* also triggers loss of TX protein, as found in *Tsn* KO mice. In contrast, conditional deletion of *Tsnax* does not affect TN protein levels. Furthermore, we confirmed that the floxed alleles of *Tsn* or *Tsnax* do not affect the basal expression levels of these proteins (Figure 5D). As these experiments validated these conditional deletion lines, we monitored the impact of conditional deletion of *Tsn* or *Tsnax* from adipocytes on adiposity. We found that conditional deletion of either *Tsn* or *Tsnax* in adipocytes in either male or female mice does not phenocopy the increased adiposity found in *Tsn* KO mice (Figures 5B, 5C, 5E and 5F).

**Figure 5:**
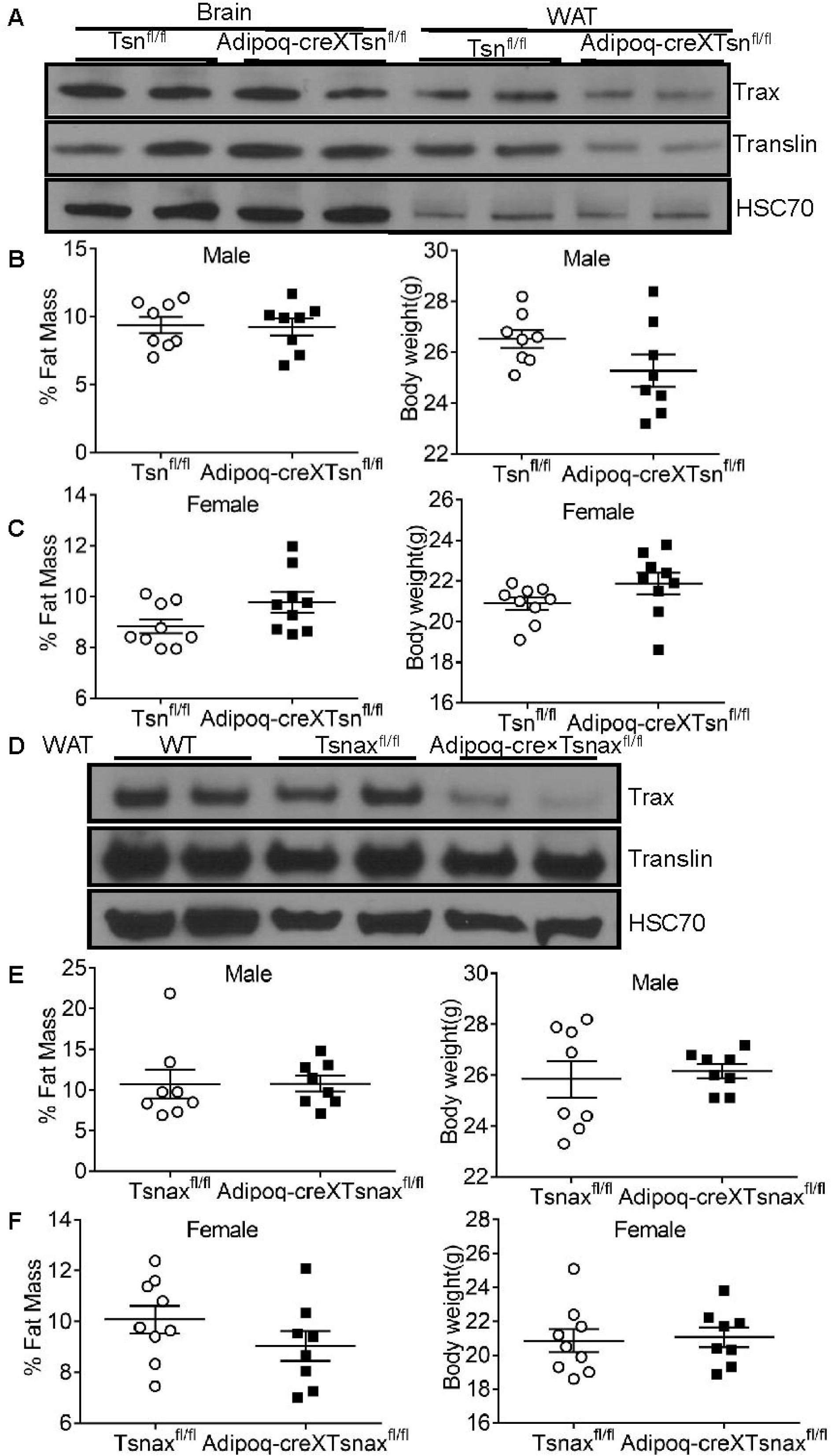
Conditional deletion of *Tsn* or *Tsnax* from adipocytes does not affect adiposity. (A) Immunoblot presented in this panel confirms selective, conditional deletion of TN protein from adipocytes in white adipose tissue (WAT) but not brain of Adipoq-Cre x Tsn^fl/fl^ mice. Note that conditional deletion of *Tsn* also produces a comparable reduction in TX protein, as found in global *Tsn* KO mice. Body weight and fat mass are normal in Adipoq-Cre xTsn^fl/fl^ male (B) and female (C) mice. (D) Immunoblot confirms conditional deletion of TX protein in Adipoq-Cre x Tsnax^fl/fl^ mice, while TN protein is unaffected. Both male (E) and female (F) Adipoq-Cre xTsnax^fl/fl^ mice have the same body weight and fat mass as Tsnax^fl/fl^ mice. n=8-9/group. Data are expressed as mean ± SEM. Student’s t-test was used for statistical analysis.

To assess if selective loss of TX from adipocytes might alter glucose tolerance in the absence of adiposity, we monitored the response of *Tsnax* conditional KO mice to a glucose challenge and found that it does not differ from that of control littermates that are homozygous for the *Tsnax*^fl/fl^ allele (Figure S5). Selective loss of TX from adipocytes also does not affect plasma levels of adiponectin in these mice (Figure S5).

### 3.4. Conditional deletion of Tsn or Tsnax from hepatocytes: effect on adiposity and transcriptional response to glucose

To check whether conditional deletion of *Tsn* or *Tsnax* from hepatocytes might phenocopy the increased adiposity exhibited by *Tsn* KO mice, we scanned mice that were homozygous for either the *Tsn* or *Tsnax* floxed allele and hemizygous for the albumin-Cre allele to drive Cre expression selectively in hepatocytes. These studies also yielded negative results (Figure 6).

**Figure 6:**
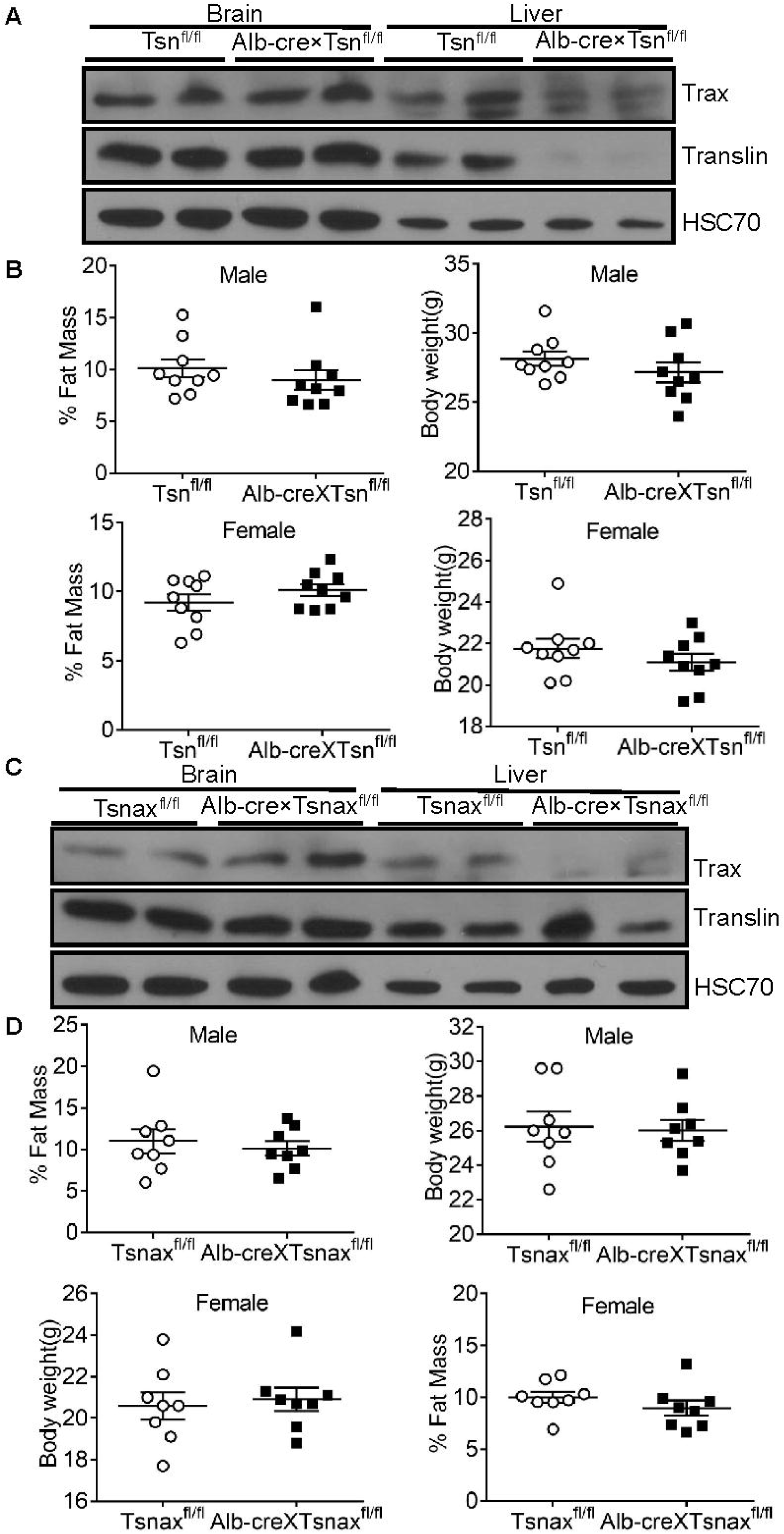
Conditional deletion of *Tsn* or *Tsnax* from hepatocytes does not affect adiposity. (A) Immunoblotting confirms selective, conditional deletion of TN from hepatocytes in Alb-Cre xTsn^fl/fl^ mice. As found in adipocytes, conditional deletion of TN produces a parallel loss of TX protein in liver. Conditional deletion of *Tsnax* from hepatocytes in either male (B) or female (C) does not affect body weight or fat mass. (D) TX protein was deleted in Alb-Cre x Tsnax^fl/fl^ mice, while TN protein levels are unaffected. Alb-Cre x Tsnax^fl/fl^ mice showed the same body weight and fat mass as Tsnax mice in both male (E) and female (F). n=8-9/group. Data are expressed as Mean ± SEM. Student’s t-test was used for statistical analysis.

Previous studies found that the levels of TN protein in hepatocytes respond to glucose levels[28, 29]. Furthermore, those studies implicated TN in transcriptional regulation of several genes that contain the glucose response element in their promoters. Since mice with a conditional deletion of *Tsn* provide an excellent opportunity to test this hypothesis, we checked whether several genes that are transcriptionally regulated by the glucose response element, LPK (Liver pyruvate kinase, FAS (Fatty acid synthase), ACC1 (Acetyl-CoA carboxylase 1) and PEPCK (Phosphoenolpyruvate carboxykinase), display altered responses to glucose in these mice. However, we did not detect any effect of conditional deletion of *Tsn* from hepatocytes on transcriptional activation of these glucose-responsive genes (Figure S6).

### 3.5. Effect of global, conditional deletion of Tsn in adulthood on adiposity

The ability of *Tsn* deletion or the *Tsnax*(E126A) point mutation to increase adiposity in the context of normal body weight suggests that it may act by biasing mesenchymal progenitor cell differentiation during development toward the adipocyte lineage and away from differentiation into osteocytes and myocytes. Furthermore, recent in vitro studies with mesenchymal stem cells support this hypothesis[30]. Relevant to this point, we have found increased adiposity as early as 6 weeks of age[5], the earliest time point measured, confirming that this phenotype appears during development. To test the possibility that deletion of *Tsn* during early development is necessary to drive the adiposity phenotype, we asked if global deletion of *Tsn* in adulthood is sufficient to elicit increased adiposity. To this end, we generated mice that are homozygous for the *Tsn*^fl/fl^ allele and hemizygous for the ubiquitin C-CreERT2 allele, which drives global expression of a Cre variant that can be activated by tamoxifen[31]. Tamoxifen was administered at 3 months of age and the mice were scanned monthly for 3 months. Although we confirmed that tamoxifen treatment is highly effective at suppressing TN (and TX) expression within 3 weeks following treatment ubiquitin C-CreERT2 mice, it did not affect body weight or adiposity in these mice (Figure 7).

**Figure 7:**
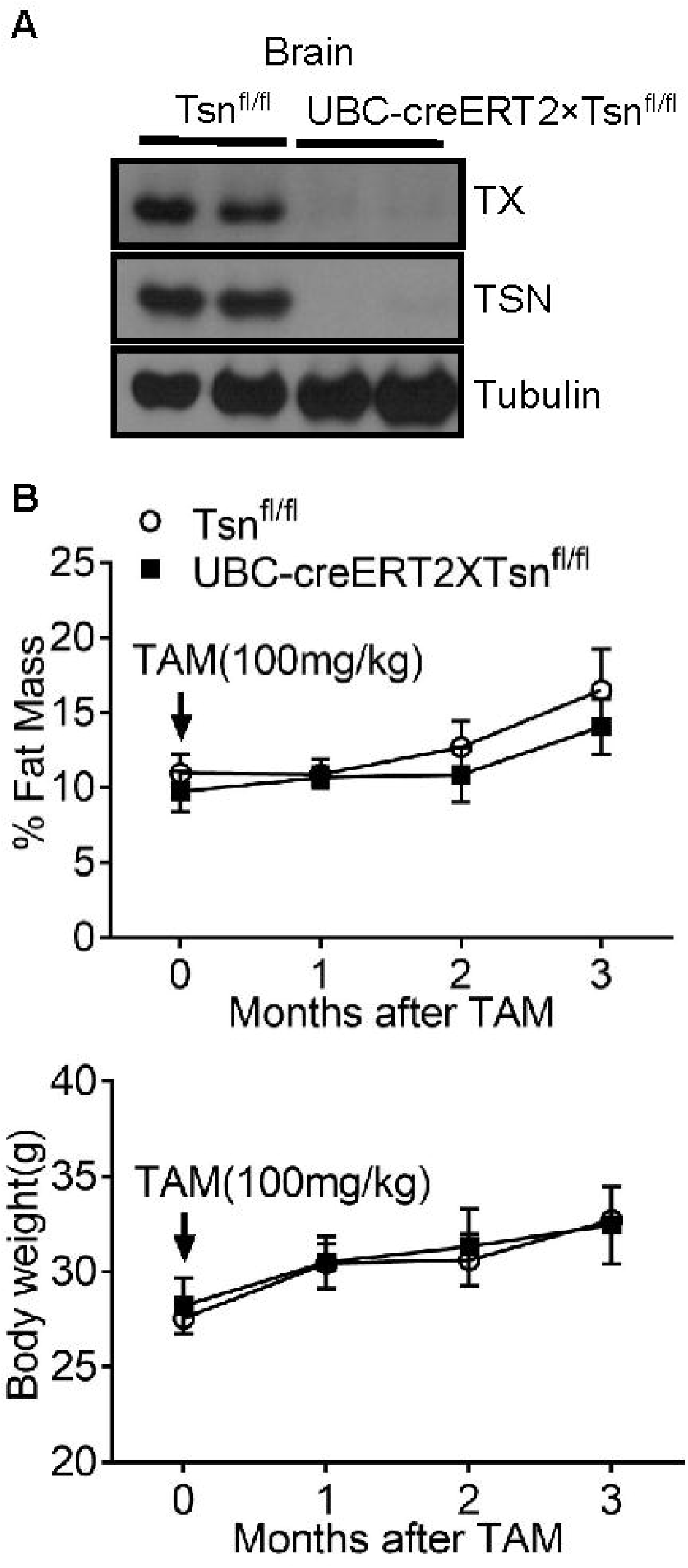
Global conditional deletion of *Tsn* in adulthood does not elicit increased adiposity. (A) Immunoblot analysis of forebrain lysates confirms that tamoxifen treatment initiated at 3 months of age produces the expected conditional deletion of *Tsn* in UBC-CreERT2 x Tsn^fl/fl^ mice, with parallel loss of TX protein. (B) UBC-creERT2 x Tsn^fl/fl^ mice have normal body weight and fat mass after tamoxifen (TAM) injection. n=6/group. Data are expressed as mean ± SEM. Two-way ANOVA with repeated measures followed by Bonferroni’s post-hoc test was used for statistical analysis.

## 4. Discussion

A top priority of current research in metabolism is to delineate the molecular mechanisms that regulate adipose tissue size and health. *Tsn* KO mice display an unusual metabolic profile, i.e. robust adiposity in the context of normal or minimally changed body weight, along with normal glucose tolerance. Therefore, we have pursued studies aimed at elucidating the molecular mechanisms linking *Tsn* deletion to increased adiposity. To this end, we have characterized the impact of genetically inactivating the TN/TX microRNA-degrading enzyme by inserting the E126A point mutation into TX. Furthermore, to gain information about which cell type(s) might drive this phenotype, as well as when TN loss is needed to induce the adiposity phenotype, we have generated and characterized the impact of conditional deletion of *Tsn* or *Tsnax* in candidate cell types and in adulthood.

We have found that mutating a single amino acid in TX, E126A, which completely inactivates its microRNA-degrading activity, is sufficient to phenocopy the robust adiposity displayed by deletion of TN, which also produces complete loss of TX protein [32]. Thus, this finding indicates that this striking phenotype is due to impaired degradation of microRNAs targeted by the TN/TX complex. Comparison of microRNA profiles in several tissues from WT and TN KO mice, or following TN knockdown in several cell lines, has identified increases in a small subpopulation of microRNAs in each tissue or cell line that only partially overlap[10, 12, 13]. Accordingly, it will be important in future studies to identify which cell types drive the adiposity phenotype, as a starting point for identifying candidate microRNAs that mediate this phenotype.

Although we have provided compelling evidence that the Tsnax(E126A) mutation abolishes the microRNA-degrading activity of the TN/TX complex, it is important to keep in mind that this mutation might also disrupt other cellular effects of TX. The observation that deletion of Tsn leads to the absence of TX protein suggested that TX is unstable unless bound to TN [8]. However, more recent studies have called this inference into question as they provide evidence that TX can act in a TN-independent fashion by binding to and activating ATM kinase in response to genotoxic stress [33]. However, these studies found that TX only associates with ATM in response to genotoxic stress leaving it unclear if TX can act in a TN-independent fashion under physiological conditions. In addition, they confirmed the TX(E126A) mutation does not impair the ability of TX to activate ATM kinase in response to genotoxic stress. Thus, it seems reasonable to attribute the ability of the Tsnax(E126A) mutation to phenocopy the robust adiposity exhibited by Tsn KO mice to its inactivation of the microRNA-degrading activity of the TN/TX complex.

Both Tsn and Tsnax are broadly expressed throughout the body (Gene Cards, Weizmann Institute). Thus, their tissue distribution provides little guidance about which cell types might drive the adipocyte phenotype. To begin the process of identifying the cell types that drive the adiposity phenotype due to loss of TN, we examined the impact of conditional deletion of *Tsn* or *Tsnax* from adipocytes to check if this phenotype might be cell autonomous. Furthermore, we also checked the impact of conditional deletion of these genes from hepatocytes, an obvious candidate given their critical role in regulating energy balance. Although neither of these manipulations elicited the adiposity phenotype, it is important to note that the adiponectin promoter appears to become active only in mature adipocytes[27]. Thus, it is conceivable that TN deletion from adipocyte or mesenchymal precursor cells may stimulate adipogenesis and contribute to the adiposity phenotype. This scenario would be consistent with the report that *Tsn* deletion increases differentiation of both mesenchymal stem cells and adipocyte precursor cells *in vitro*[30].

In our prior characterization of the adiposity phenotype in *Tsn* KO, we noted that elevated adiposity is present at 6 weeks of age, the earliest time point assayed, and continues to increase through 10 months of age, the oldest time point checked. Thus, it seemed reasonable to assume that the absence of TN in adulthood would be sufficient to induce increased adiposity. To test this hypothesis, we examined the impact of inducing global, conditional deletion of TN at 3 months of age and were surprised to find that this manipulation does not increase adiposity. Thus, this result implies that deletion of *Tsn* during development is needed to drive the adiposity phenotype.

Epigenetic modifiers, including miRNAs[34], have an important role in developmental programming and impart long lasting phenotypic changes. Proper hypothalamic development, for example, is dependent on metabolic hormones, such as leptin, ghrelin, and insulin[35, 36], that act as neurotrophic factors during the neonatal period and do not exert their actions related to energy balance until the adolescent/adult ages. It is critical that these hormones are secreted in the appropriate pattern (developmental timing, concentration) to facilitate proper brain development and control of energy balance[37]. Administration of leptin, for example, to a leptin deficient *Ob/Ob* mouse during the neonatal period rescues deficient hypothalamic neural development, but will not have an effect if administered in adulthood indicating a critical sensitive developmental time window[38]. In contrast, elevated leptin levels during the neonatal period are also detrimental to the developing brain and lead to hypothalamic leptin resistance and obesity[39]. While these neurotrophic effects have been well-studied in the hypothalamus, other brain regions, as well as other organ systems, may also be subject to such developmental programming influences that have long-term consequences through adulthood.

Our observation that global, conditional deletion of *Tsn* in early adulthood does not elicit adiposity has important practical implications. In recent studies we have found that *Tsn* deletion confers protection against developing pathogenic vascular stiffness[13]. Those results indicate that compounds that inhibit the TN/TX complex may have therapeutic potential in preventing larger artery stiffness, a major public health problem[40]. Our finding that *Tsn* deletion during adulthood, which would be similar to systemic administration of an inhibitor of TN/TX RNase activity, does not elicit increased adiposity enhances the translational potential of this therapeutic strategy.

In this study, we have used *Tsnax*(E126A) mice to assess if the metabolic profile displayed by *Tsn* KO mice can be attributed to inactivation of the TN/TX microRNA-degrading enzyme, which leads to increased levels of a small population of microRNAs, in a cell-type specific fashion. Remarkably, *Tsnax*(E126A) and *Tsn* KO mice display nearly identical adiposity. Furthermore, in the course of more extensive metabolic profiling of these mice, we did not detect significant differences between them. However, it is worth noting that in many cases, such as GTT assays, *Tsnax*(E126A) mice are significantly different from WT mice, while *Tsn* KO mice are not. Thus, it is conceivable that these subtle differences may reflect additional cellular effects caused by loss of TN and TX proteins in *Tsn* KO mice, in addition to inactivation of the TN/TX microRNA-degrading enzyme. From this perspective, it will be interesting to check whether other phenotypes exhibited by *Tsn* KO mice, such as blockade of pathogenic vascular stiffness[13] and defects in synaptic plasticity[12] are mimicked by *Tsnax*(E126A) mice.

## 5. Conclusion

Mice containing a point mutation in Tsnax, E126A, that inactivates the TN/TX microRNA-degrading enzyme phenocopy the robust adiposity displayed by Tsn KO mice. Furthermore, global conditional deletion of Tsn in adulthood does not elicit increased adiposity. Taken together, these findings indicate that inactivation of the TN/TX RNase during development drives the robust adiposity displayed by Tsn KO mice.

## Acknowledgments

This study was supported by funds from NIDA (P50 DA044123; JMB) and the Mid-Atlantic Nutrition and Obesity Research Center (JMB). We thank Dr. Z. Paroo for providing recombinant TN/TX proteins, Dr. Michael Wolfgang for helpful discussions and the Hopkins Transgenic Core for expert assistance in generating mouse lines.

## Abbreviations

Tsn: Translin
Tsnax: Translin associated protein X
TN: Translin
TX: Trax

## Supplementary Figures and Tables

**Figure S1:**
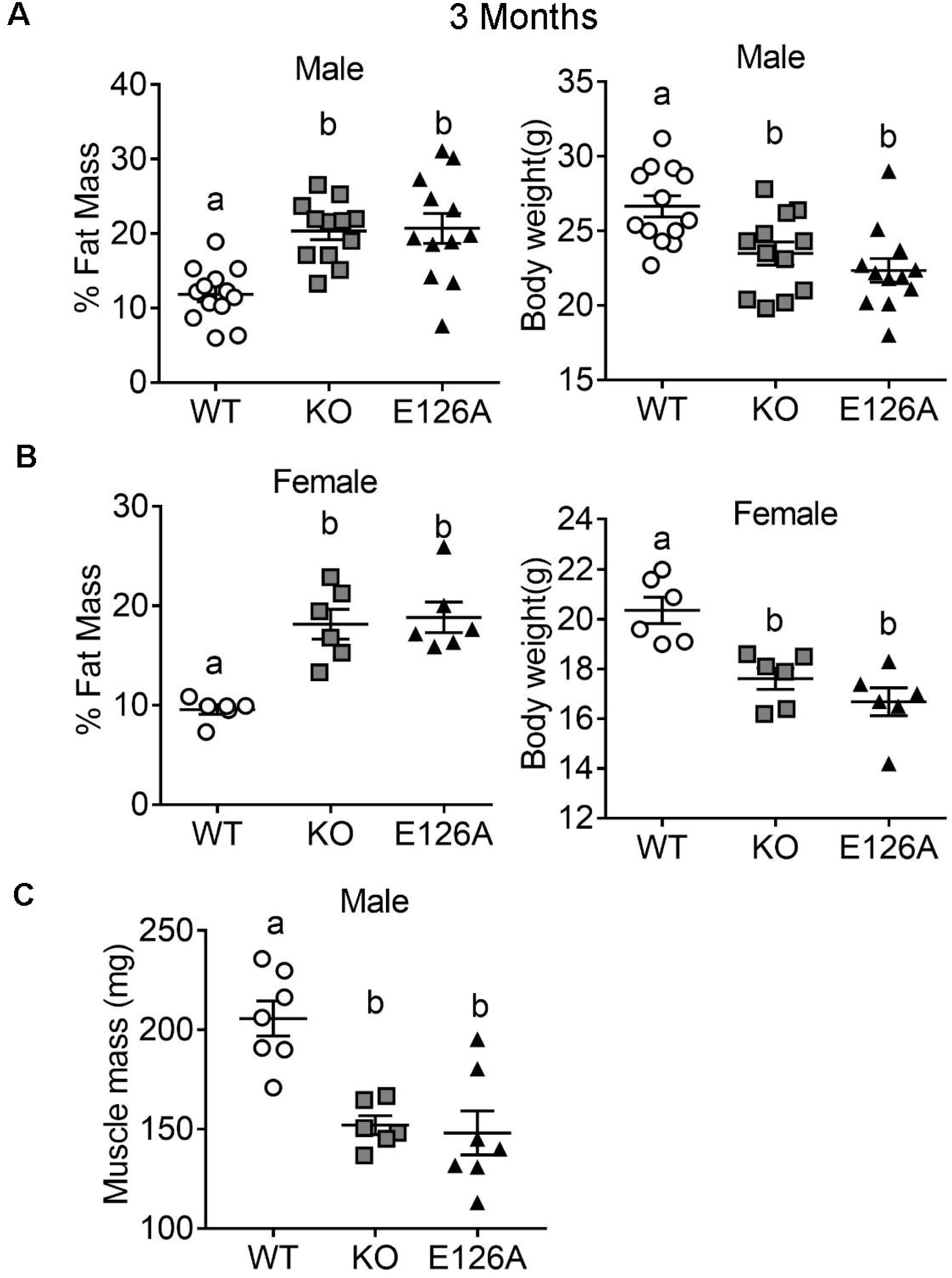
*Tsnax*(E126A) mice phenocopy increased adiposity and decreased muscle mass displayed by *Tsn*. KO mice (A and B) At 3 months of age, both male and female Tsnax(E126A) mice match the increased adiposity and decreased body weight displayed by *Tsn* KO mice. (C) Both *Tsn* KO mice and *Tsnax*(E126A) mice show reduced muscle mass as determined by weighing the quadriceps femoris muscle. n=6-13/group. Data are expressed as mean ± SEM. Different letters indicate statistically significant differences (P < 0.05) by one-way ANOVA followed by Bonferroni’s post-hoc test.

**Figure S2:**
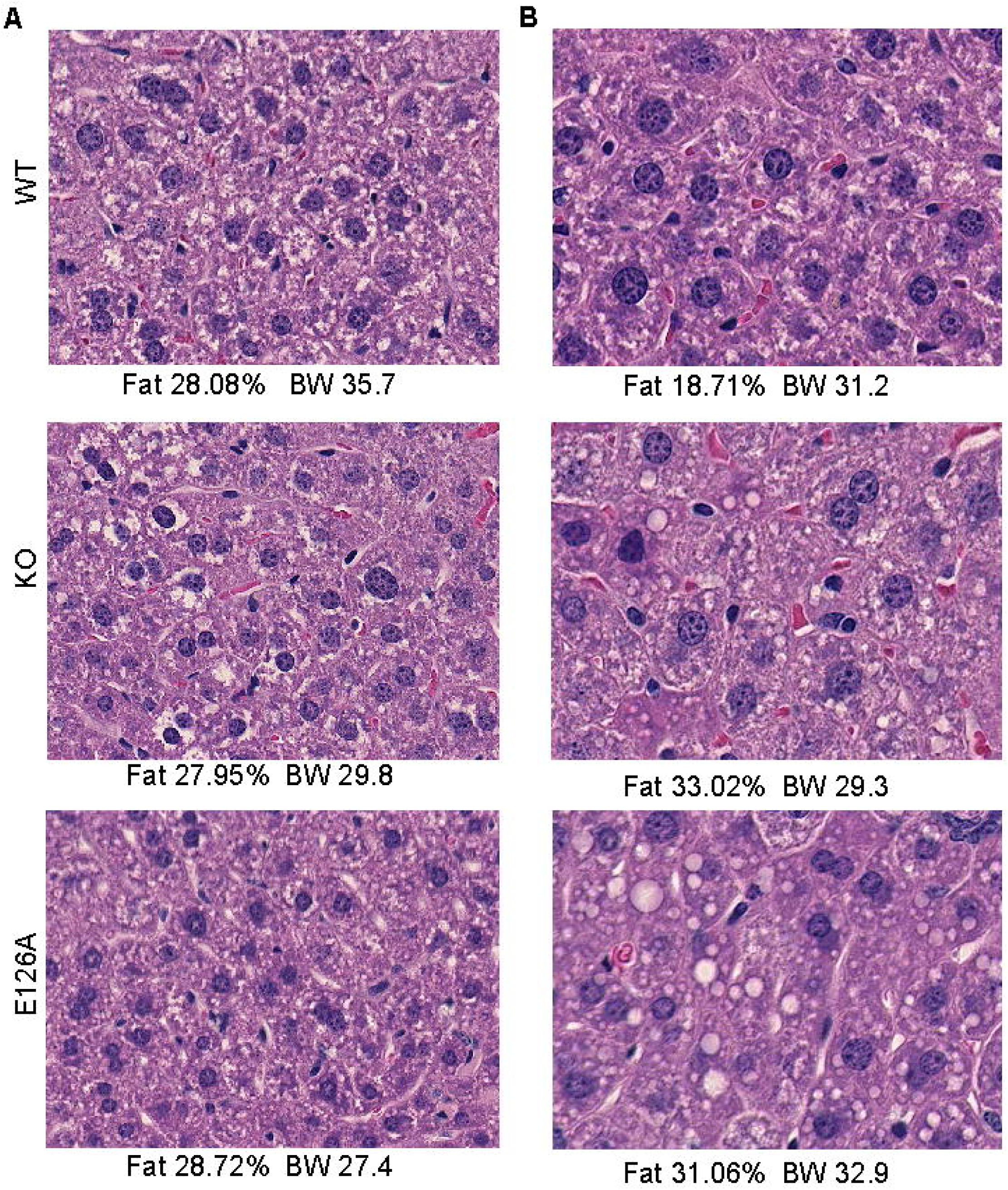
Representative images of H&E stained liver sections from WT, *Tsn* KO and *Tsnax*(E126A) mice. Lipid droplets are present in sections taken from male *Tsn* KO and *Tsnax*(E126A) mice that have the highest levels of adiposity and body weight (B), but not in sections taken from *Tsn* KO and *Tsnax*(E126A) mice with lower levels of adiposity and body weight (A).

**Figure S3:**
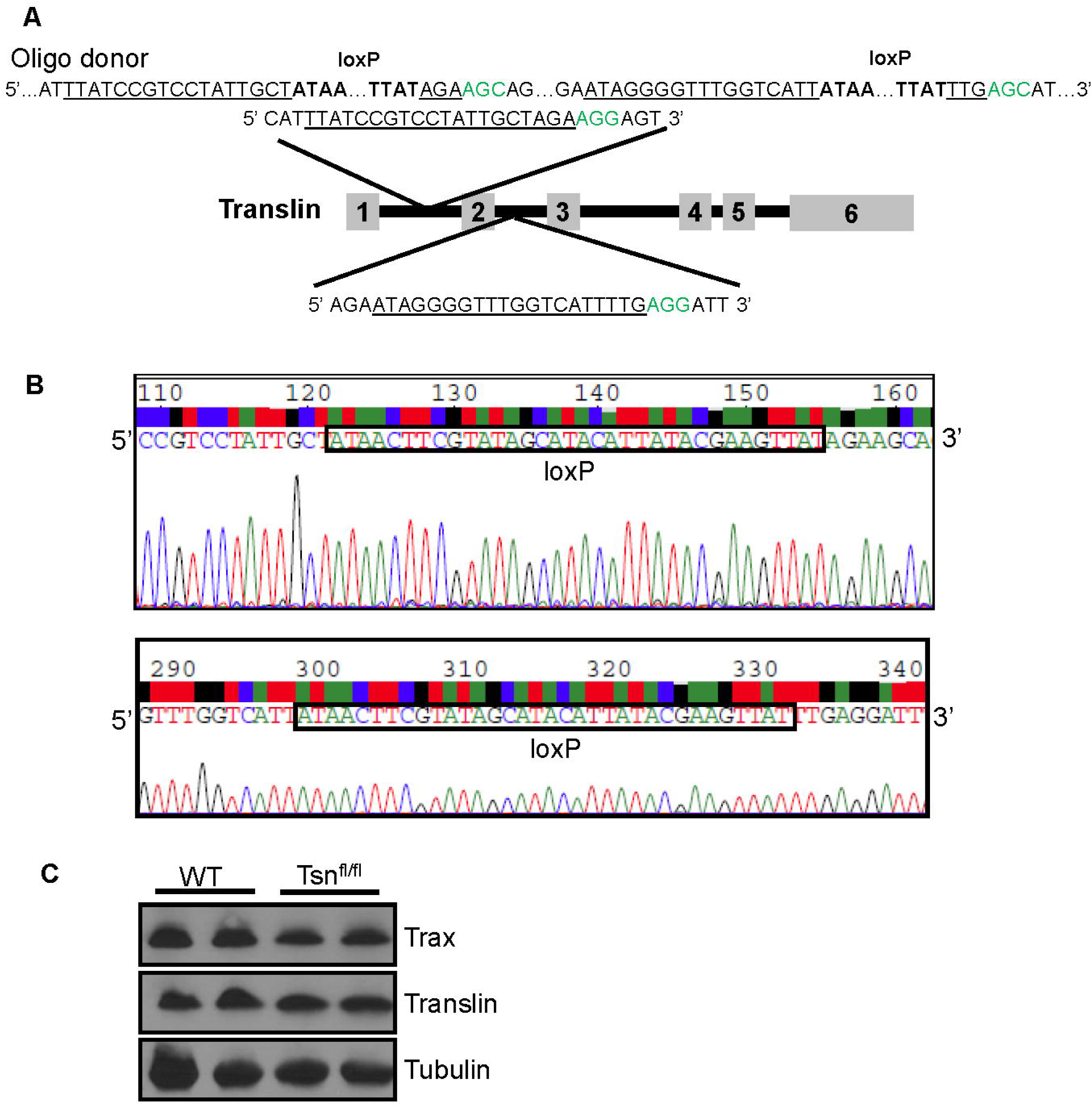
Generation of mice with a floxed *Tsn* allele. (A) Schematic representation of the *Tsn* gene segment matching the designed sgRNA and oligo donor used for inserting loxP sites by CRISPR. PAM site is marked by green. Underlined segments show the location of the guide RNA sequences. (B) The nucleotide sequence of the segments matching sgRNA sequences in Tsn^fl/fl^ mice demonstrates the successful insertion of loxP sites. (C) Immunoblotting shows that TN and TX protein expression levels are normal in forebrain of Tsn^fl/fl^ mice.

**Figure S4:**
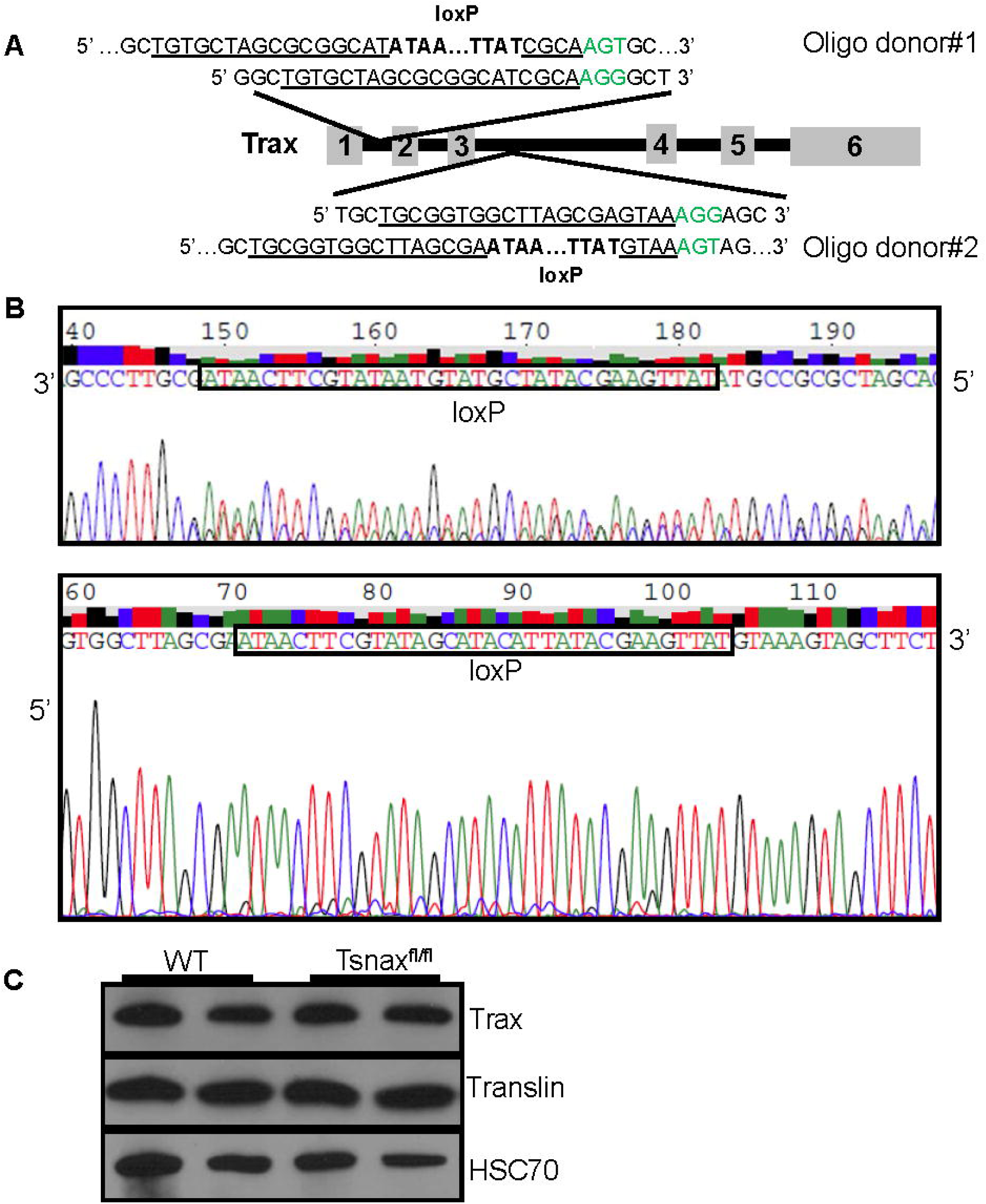
Generation of mice with a floxed *Tsnax* allele. (A) Schematic representation of the *Tsnax* gene region matching the designed sgRNA and oligo donors used for inserting loxP sites by CRISPR. PAM site is marked by green. Underlined segments show the positions of the guide RNA sequences. (B) The nucleotide sequences of the sgRNA matching regions of Tsnax^fl/fl^ mice confirm insertion of loxP sites at the intended locations. (C) Immunoblotting of forebrain samples confirms that TN and TX protein levels are normal in forebrain of Tsnax^fl/fl^ mice.

**Figure S5:**
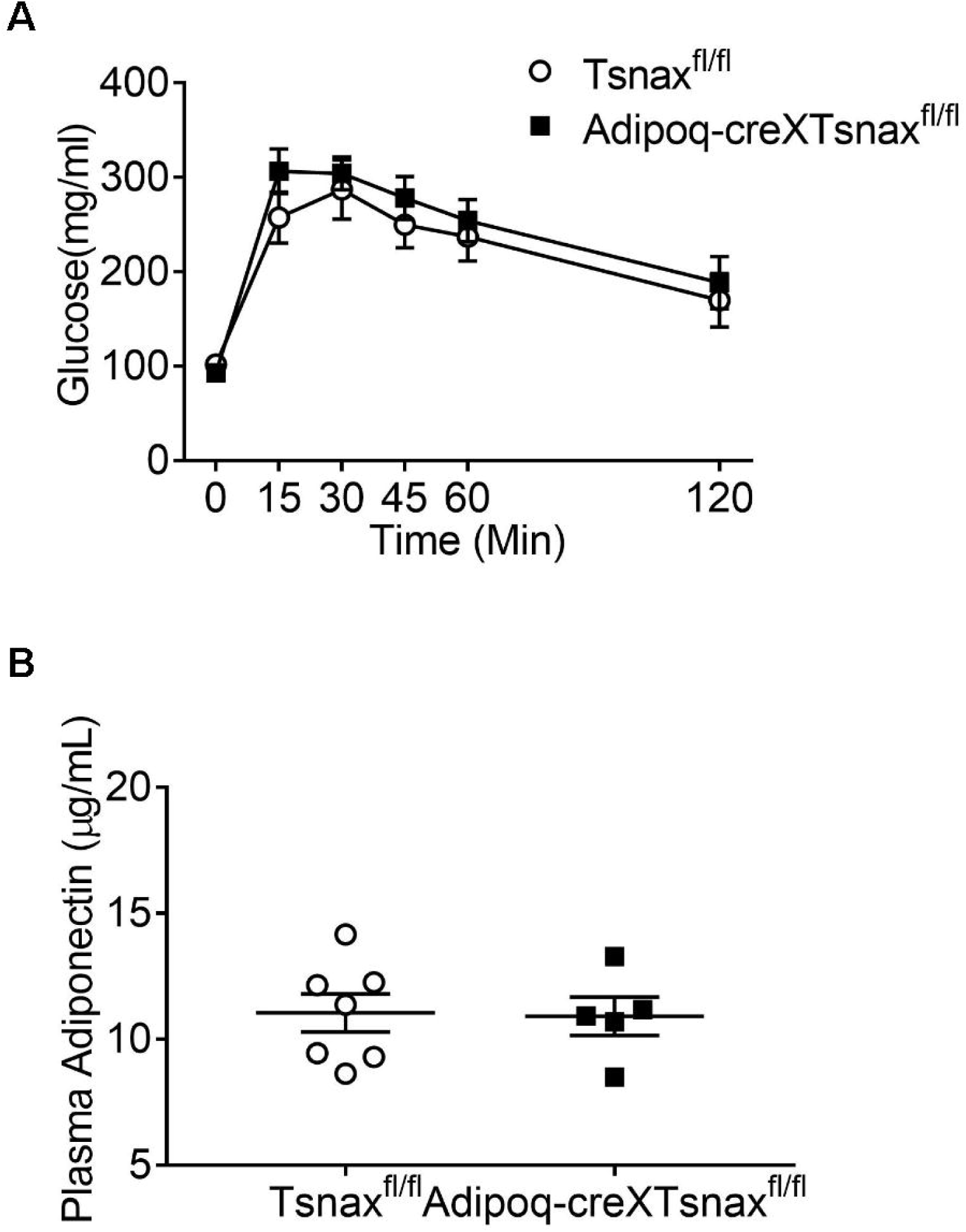
Normal glucose tolerance and plasma adiponectin levels in Adipoq-Cre x Tsnax^fl/fl^ mice. Blood glucose levels during ip-GTT (1.5 mg/g glucose) of Adipoq-Cre xTsnax^fl/fl^ male mice are normal compared with Tsnax^fl/fl^ littermates. (B) Plasma adiponectin levels are unaffected by conditional deletion of *Tsnax* from adipocytes. n=6/group. Data are expressed as mean ± SEM. Two-way ANOVA with repeated measures followed by Bonferroni’s post-hoc test or Student’s t-test were used for statistical analysis.

**Figure S6:**
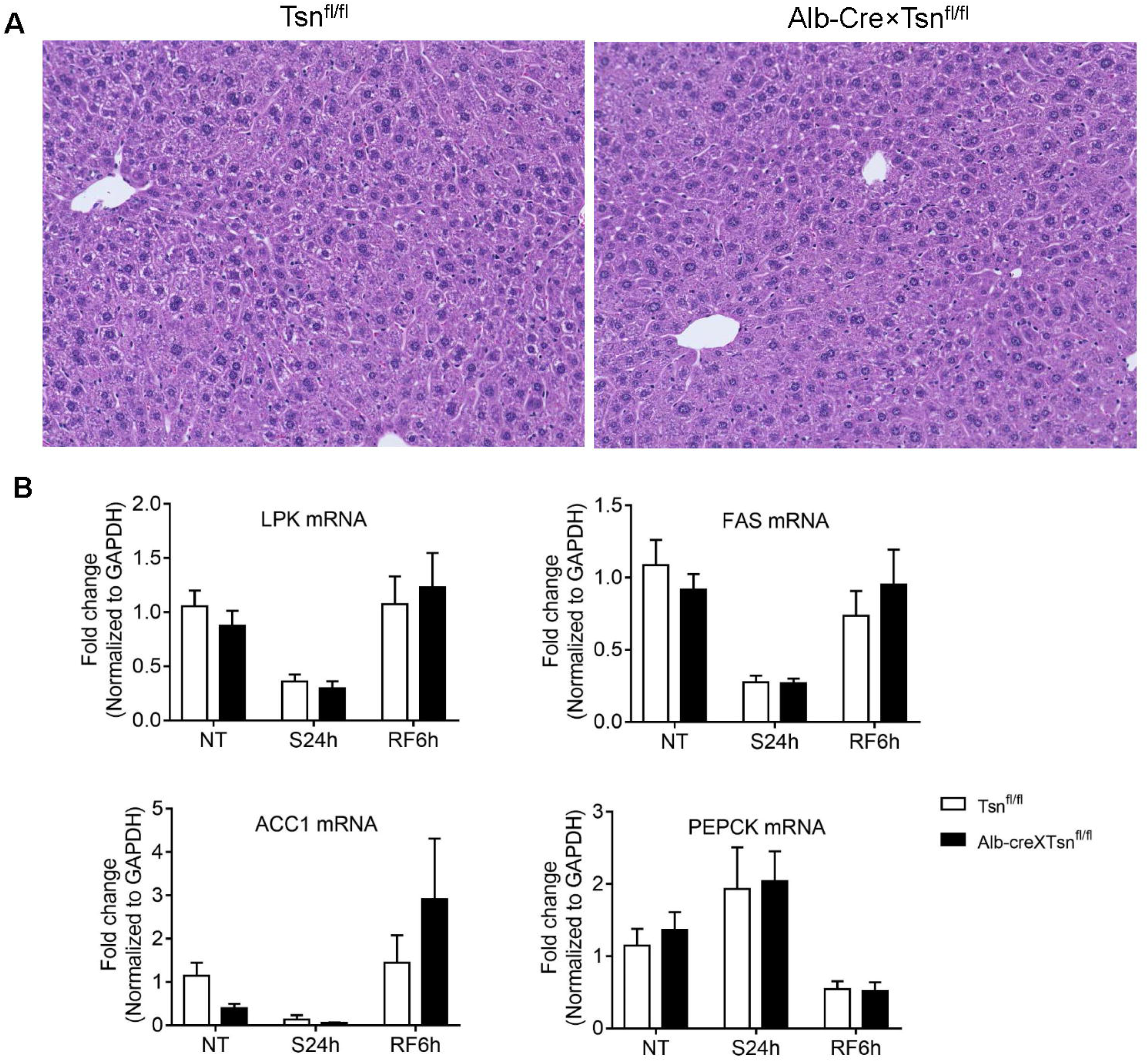
Normal transcriptional response to glucose in Alb-Cre xTsn^fl/fl^ male mice. H&E stained liver sections taken from Alb-Cre xTsn^fl/fl^ and Tsn^fl/fl^ mice show normal histology. (B) RT-PCR of Alb-Cre xTsn^fl/fl^ liver tissue shows normal mRNA levels of LPK, ACC1, FAS, PEPCK. NT: no treatment. S24h: starvation for 24 h. RF6h: re-fed with high sucrose diet for 6 h. n=6/group. Data are expressed as mean ± SEM. Two-way ANOVA with repeated measures followed by Bonferroni’s post-hoc test was used for statistical analysis.

**Supplementary Table 1.**
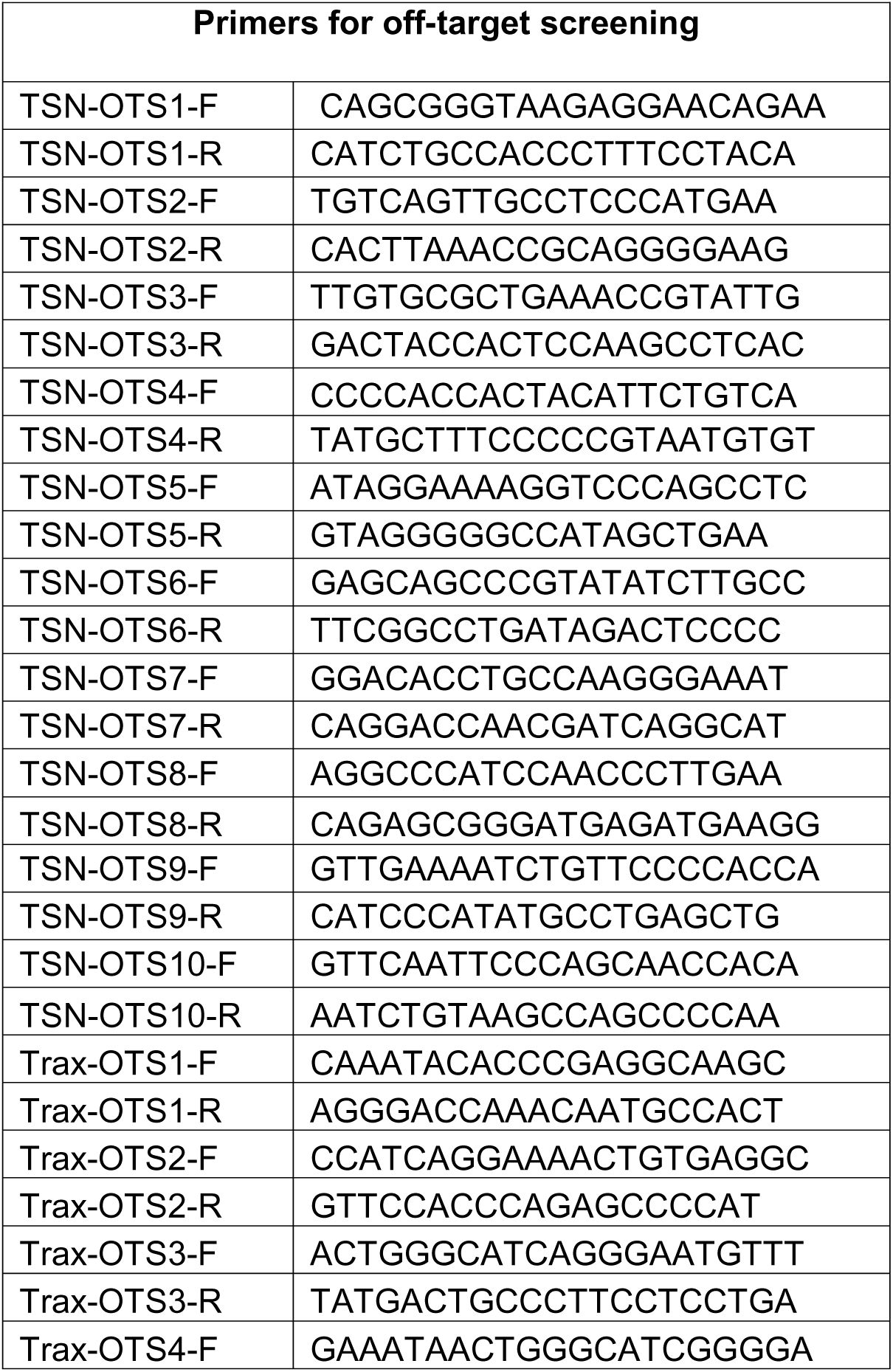
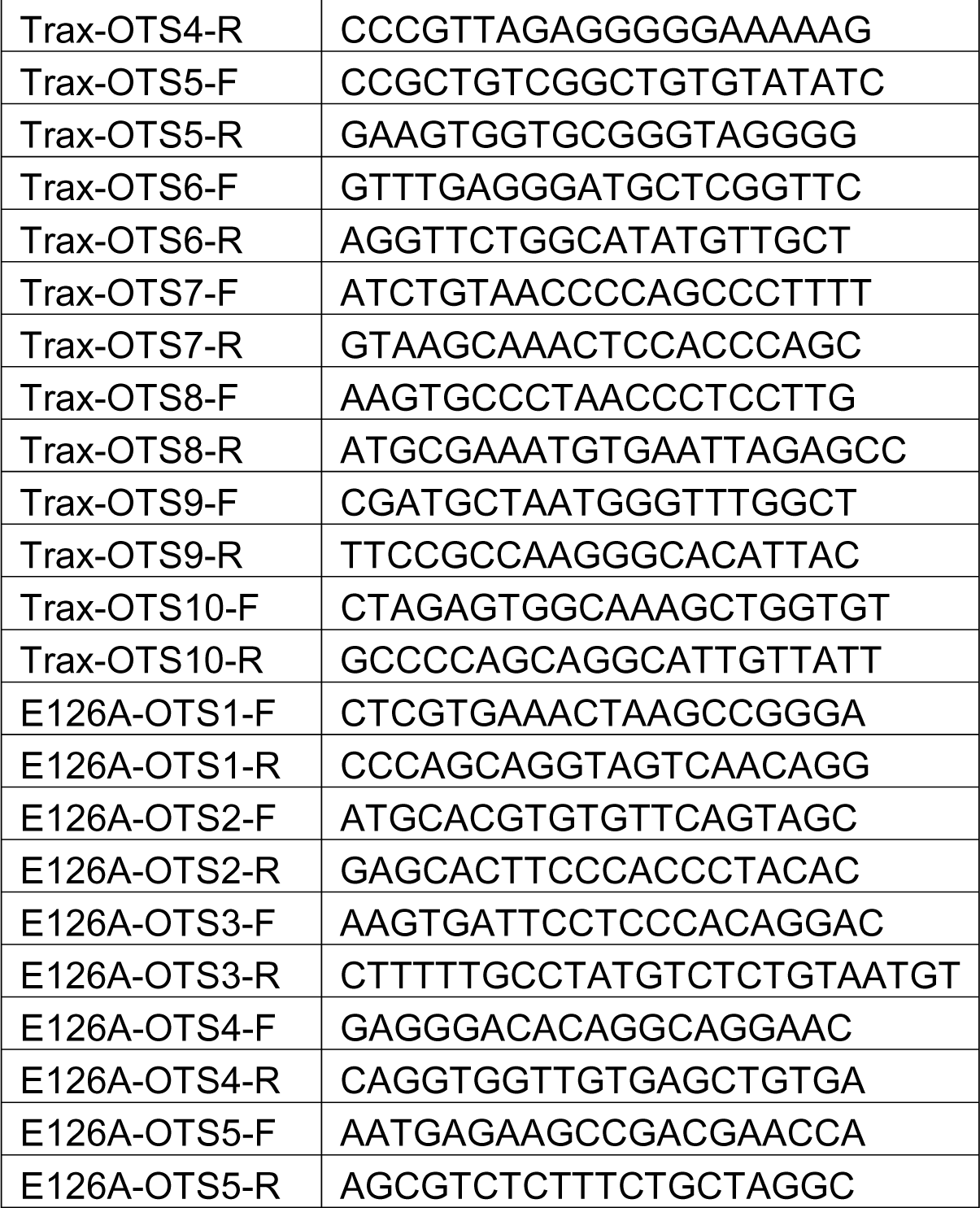

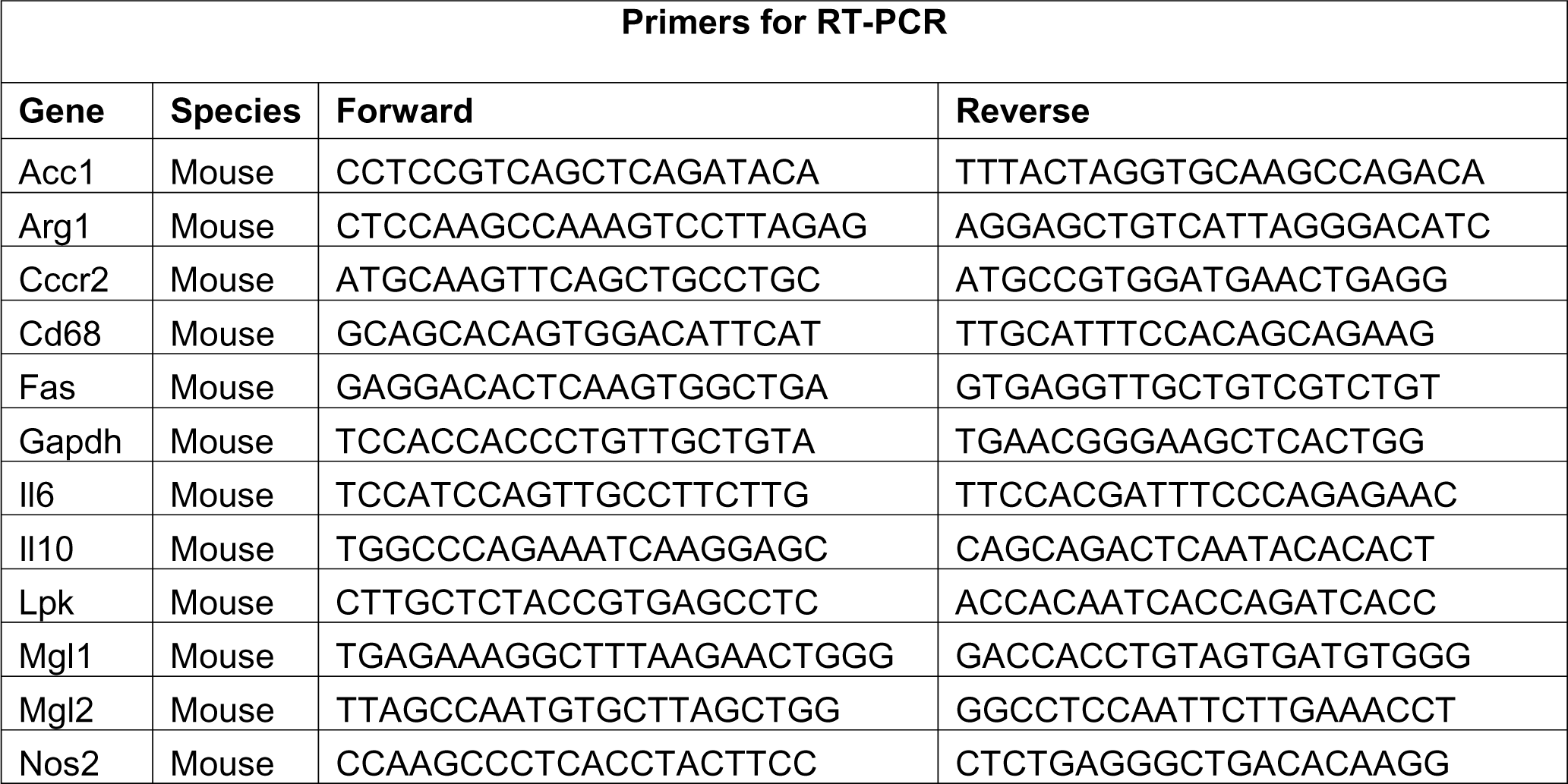
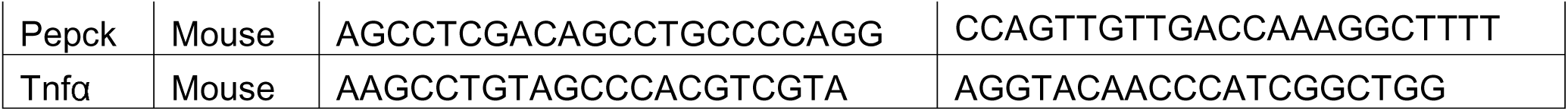

